# A scalable, data analytics workflow for image-based morphological profiles

**DOI:** 10.1101/2023.07.03.547611

**Authors:** Edvin Forsgren, Olivier Cloarec, Pär Jonsson, Johan Trygg

**Affiliations:** Umeå University CLiC, Department of Chemistry, Umeå, Sweden; Sartorius Stedim Data Analytics, Corporate Research, Umeå, Sweden

## Abstract

Cell Painting is an established community-based, microscopy-assay platform that provides high-throughput, high-content data for biological readouts. In November 2022, the JUMP-Cell Painting Consortium released the largest annotated, publicly available dataset, comprising more than 2 billion cell images. This dataset is designed for predicting the activity and toxicity of 100k drug compounds, with the aim to make cell images as computable as genomes and transcriptomes.

In this paper, we have developed a data analytics workflow that is both scalable and computationally efficient, while providing significant, biologically relevant insights for biologists estimating and comparing the effects of different drug treatments.

The two main objectives proposed include: 1) a simple, yet sophisticated, scalable data analytics metric that utilizes negative controls for comparing morphological cell profiles. We call this metric the equivalence score (Eq. score). 2) A workflow to identify and amplify subtle morphological image profile changes caused by drug treatments, compared to the negative controls. In summary, we provide a data analytics workflow to assist biologists in interpreting high-dimensional image features, not necessarily limited to morphological ones. This enhances the efficiency of drug candidate screening, thereby streamlining the drug development process. By increasing our understanding of using complex image-based data, we can decrease the cost and time to develop new, life-saving treatments.

**Author summary:** Microscopy-assays are often used to study cell responses to treatments in the search for new drugs. In this paper, we present a method that simplifies the understanding of the data generated from such assays. The data in this study consists of 750 morphological features, which describe the traits and characteristics of the cells, extracted from the images. By using untreated cells as a biological baseline, we’re able to detect subtle changes that occur in the treated cells. These changes are then transformed into an equivalence score (Eq. score), a metric that lets us compare the similarities among different treatments relative to our baseline of untreated cells. Our Eq. score approach transforms complex, high-dimensional data about cell morphology into something more interpretable and understandable. It reduces the “noise” in the features and highlights important changes, the “signal”. Our method can be integrated into existing workflows, aiding researchers in understanding and interpreting complex morphological data derived from cell images more easily. Understanding cell morphology is crucial to deepening our knowledge of biological systems. Ultimately, this could contribute to the faster and more cost-effective development of new, life-saving treatments.

## Introduction

The process of drug development is a costly and time-consuming endeavor, prompting researchers to explore new, more efficient methods of drug candidate screening. One emerging approach is image-based profiling [1], which utilizes fluorescent markers with techniques such as Cell Painting and CellProfiler, as well as Artificial Intelligence (AI), to extract morphological features quickly and cost-effectively. CellProfiler is an open-source image analysis software tailored for high-throughput biological experiments. Developed by the Broad Institute, it enables customizable analysis pipelines and incorporates various image processing algorithms. Its flexibility allows for diverse applications in biological and biomedical research. By automating complex image analysis tasks, CellProfiler facilitates the extraction of meaningful, reproducible quantitative data, advancing high-content screening and biological image analysis [2]. The affordability and high-throughput nature of image-based profiling, in combination with providing both temporal and spatial information, make it an increasingly common choice for drug screening purposes. However, while these techniques have evolved greatly in recent years, connecting the thousands of images and their extracted features back to biology in understandable metrics remains a significant challenge.

To address this issue, many researchers are turning to AI to find a solution. AI has shown remarkable success in various applications, such as segmenting nuclei [3–5] and cell bodies [6, 7], image restoration [8, 9], image super-resolution [10, 11], and speeding up fluorescent 3D sample imaging [12]. These examples demonstrate how AI-based methods have excelled over traditional ones. Despite this success, the lack of explainability and interpretability remains a major concern, particularly when classifying and quantifying treatment data. This is partly due to the structure of the data, as most datasets have numerous controls but a limited number of replicates for the actual treatments. Combining this with a high-dimensional feature vector creates a skewed relationship between observations and features, known as “the curse of dimensionality” [13].

In addition to AI-based solutions, traditional correlation-based metrics such as cosine similarity, Pearson’s correlation, Spearman’s rank correlation, and Kendall’s rank are often used today to examine the similarity between treatments [14, 15]. While these metrics provide an overview of the data, they consider all features as equally important, making it difficult to capture and identify the unique, subtle morphological changes that separate different treatments from each other.

The skewed relationship between observations and features has been the standard in chemometric problems for decades, where multivariate analysis (MVA) has shown great success in capturing subtle differences between groups in high-dimensional data [16–18]. Therefore, we have developed a data analytics workflow that offers valuable insights for biologists in estimating and comparing the effect of different treatments. In this method, the negative control group serves as a biological baseline for predictive models. The models focus on the differences in the morphological profiles between treatments and the negative controls to identify the unique biological impact of a specific treatment. The two main objectives of this method are: 1) to create a simple, yet sophisticated, scalable metric for comparing morphological profiles, which we call the equivalence score (Eq. score). The Eq. score is a multivariate prediction of a model that has been trained to identify the relationship between a reference treatment and negative controls; 2) to identify and amplify the subtle morphological profile changes caused by a treatment compared to the negative controls. By transforming the morphological features to predicted Eq. scores of several reference compounds, we demonstrate that we can reduce noise and enhance the signal that exists in the treatments compared to the control group. With this, we provide a computational workflow based on MVA to assist biologists in interpreting high-dimensional features, not necessarily limited to morphological ones, and enhancing the efficiency of drug candidate screening, thereby streamlining the drug development process.

## Materials and methods

Correlation-based methods are widely used to identify similarities between morphological feature profiles. While these methods provide a quick and easy overview of the data, quantifying the nuances and subtle differences and similarities between treatments can be challenging. In this paper, we present a semi-supervised method for comparing the effects of treatments on cell cultures, which relies on negative controls (Fig. 1). To reduce noise and highlight structured variation within treatment groups, we compute Equivalence scores (Eq. scores) for each treatment using a PLS/OPLS model trained on a reference treatment and the negative controls. Eq. scores offer a more sophisticated metric compared to traditional correlation-based ones. By comparing Eq. scores, we can identify treatments with similar effects. Furthermore, we create Eq. scores for all 303 compounds in the dataset and use them as new features. We benchmark these Eq. scores features versus the original CellProfiler features and show an improved performance, even though the Eq. scores are based purely on the CellProfiler features.

**Fig 1.**
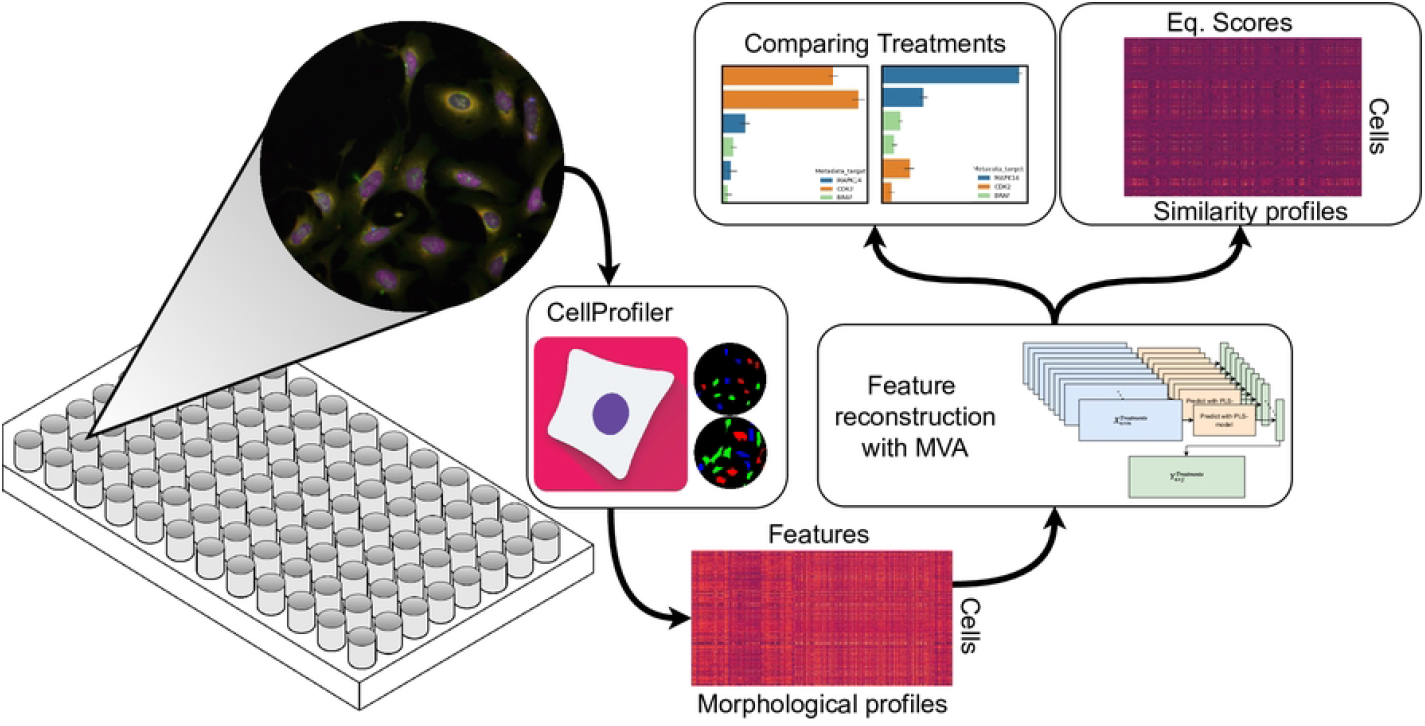
Pipeline where cells are labeled with fluorescent markers, grown in wells and then imaged through fluorescent microscopy. The images are then preprocessed and analyzed through the software CellProfiler. From the software, we get numerous features which are then transformed through our multivariate approach.

## Multivariate details

### Principal Component Analysis (PCA) -Data overview

Principal Component Analysis (PCA) is a widely-used statistical method that can be used to reduce the dimensionality of high-dimensional data. By identifying the directions of maximum variation in the data, PCA creates new, orthogonal variables called principal components. These principal components are linear combinations of the original variables and can be used to summarize the information contained in the data. In many cases, the first few principal components can capture most of the variation in the original data, allowing for easier interpretation and visualization of the data. The principal components, consisting of scores *T* and loadings *P*, are good “summaries” of *X* such as:

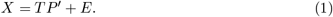

Where *E* is the residual and is “small” if enough principal components are used and/or *X* contains mainly systematic variation and low levels of noise.

### PLS/OPLS -Predictive modeling

(Orthogonal) Partial Least Squares (PLS/OPLS) regression are multivariate techniques for modeling the relationship between two matrices of variables: a predictor matrix X and a response matrix Y. Like PCA, PLS/OPLS aims to identify new latent variables that can summarize the systematic variation in the data. However, unlike PCA, PLS/OPLS takes into account the additional information provided by the response matrix Y, allowing it to model the relationship between X and Y directly.

In PLS/OPLS, the predictor matrix X is decomposed into a set of scores *T* and loadings *P*, as

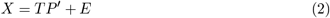

where *E* is the X-residual. In addition to this, the response matrix Y is also decomposed into a set of scores C and loadings T:

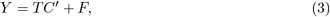

where *C* is the weights associated with *Y* and *F* is the Y-residuals. *F* represents the difference between the observed *Y* and the modeled *Ŷ* . As with PCA, the weights, scores and loadings can be calculated using various methods, with the NIPALS algorithm [19] being one of the most commonly used.

### Workflow

The premise of our approach is to compare a set of reference treatments to a control group, which serves as a baseline during a calibration phase (Fig. 2a). During the calibration phase, a scale between 0 and 1 is defined. The new scale, which we call the Eq. score, represents the proportion of equivalence of a new treatment compared to the reference treatment. PLS/OPLS regression is used to build the Eq. scores. The features corresponding to the controls and a given reference treatment (X) are regressed against an arbitrary vector of 0 and 1 (Y), corresponding to controls and the reference treatment, respectively (Fig. 2a). To ensure that the predictions of the reference group are fair, a leave-one-out cross-validation approach (LOOCV) is used for each replicate in the reference group. New treatments can then be measured in the new referential built with PLS/OPLS models from the known treatments (Fig. 2b). They can then be characterized by a single, or a spectrum, of Eq. scores and compared using multivariate data analysis.

**Fig 2.**
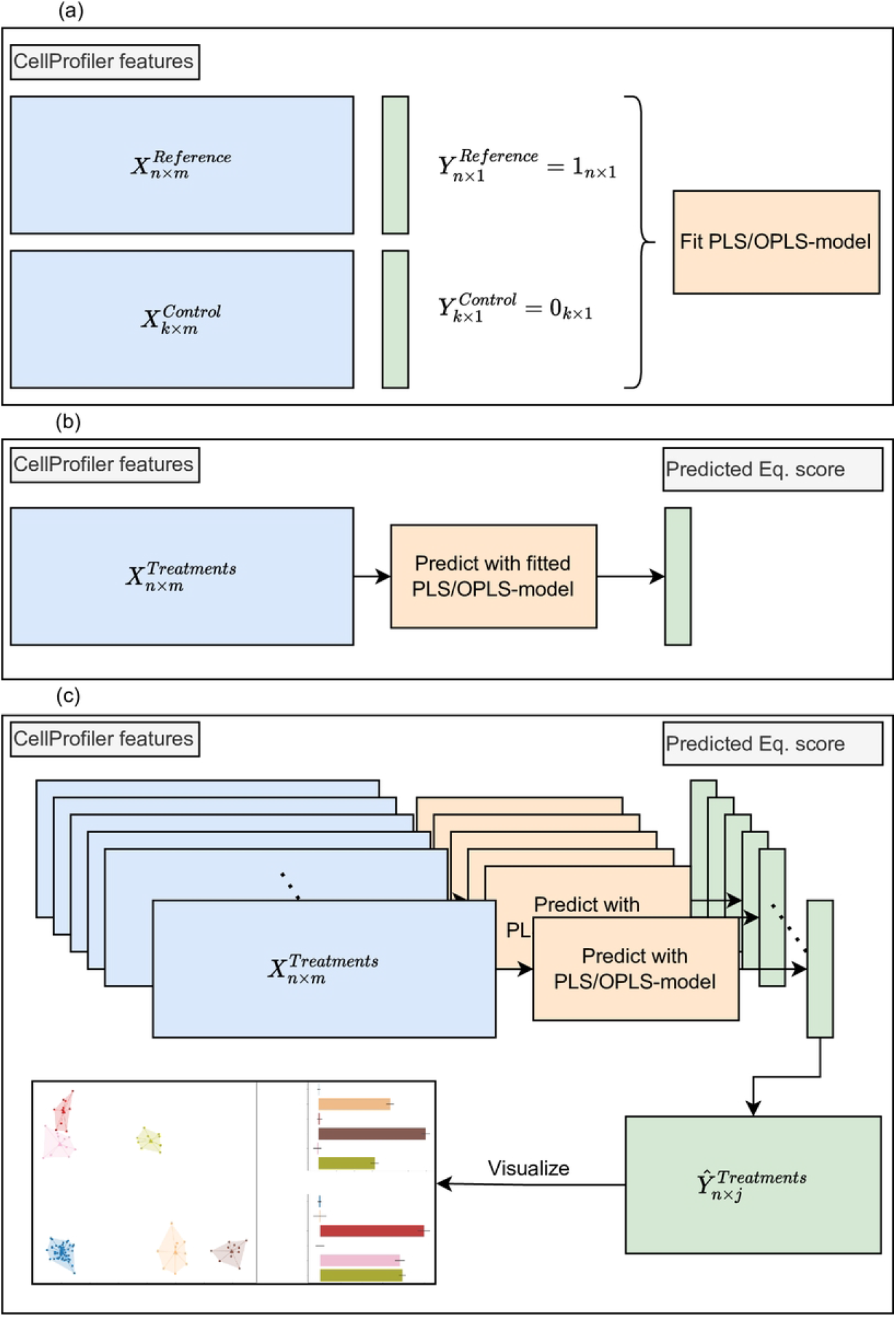
In (a) is the first step of the method, fitting a PLS/OPLS model on a reference treatment vs the control group. We create two Y-vectors where the reference observations have 1 and the control group 0. In (b) the fitted model is used on other treatments to model their Y in the same space, we call this value the Eq. score. In (c) we iterate through this process to create a new feature space consisting of the Eq. scores which can then be visualized for interpretation.

In this prediction step, the Sum Squared Error statistic (SSE) is used to evaluate the feature space (X) and the prediction error of (Y). The former indicates if the features presented by the predicted treatment lie within the training set feature space; the latter shows how well the prediction is. The SSE for observation *i* is calculated as

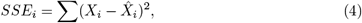

over the features and where 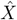 is the input *X* multiplied by the loadings and weights of the PLS/OPLS model. This means that if treatments have high Eq. scores but also high SSEs, we should be careful about drawing conclusions about similarities.

In Fig. 2c, steps (a) and (b) are repeated for each of the 303 treatments in the dataset, creating a new feature space of Eq. scores where the systematic variation is amplified and noise is reduced. This feature space, which is fully based on CellProfiler features, can then be used to look for clustering and similarities with methods such as PCA as well.

### Exemplifying with toxicity

The Eq. score method allows for the comparison of treatment effects relative to a reference treatment based on the similarity or dissimilarity of their feature profiles. One possible application of this method is in the assessment of different toxicities with different cell death mechanisms. In this section, we will use a toy dataset to demonstrate how Eq. scores can be utilized to better understand and compare the toxic effects of various treatments.

Assume we have two known toxicities, Toxicity 1 and Toxicity 2, where each serves as a reference treatment in two separate PLS/OPLS models. We can then predict the Eq. scores of other treatments i.e. Treatment 1, 2 and 3 with the two models. The Eq. scores can then be used as axes in a scatter plot to visualize the relationships among different treatments and their toxicities. Fig. 3 provide insights into the extent to which each treatment’s effects resemble the reference toxicities. From Fig. 3a, we can infer that Treatment 1 exhibits a stronger effect related to Toxicity 1, while Treatment 2 has a weaker effect related to Toxicity 2. It is possible to speculate that both treatments are the same as Toxicity 1 and 2, respectively, but at different concentrations. Interestingly, Treatment 3 appears to exhibit a combined effect of both Toxicity 1 and Toxicity 2. The same thing can be observed in Fig. 3b and c but in a bar plot for the equivalence of Toxicity 1 and 2 respectively. This type of plot is particularly useful if we only have one reference treatment we want insights into.

**Fig 3.**
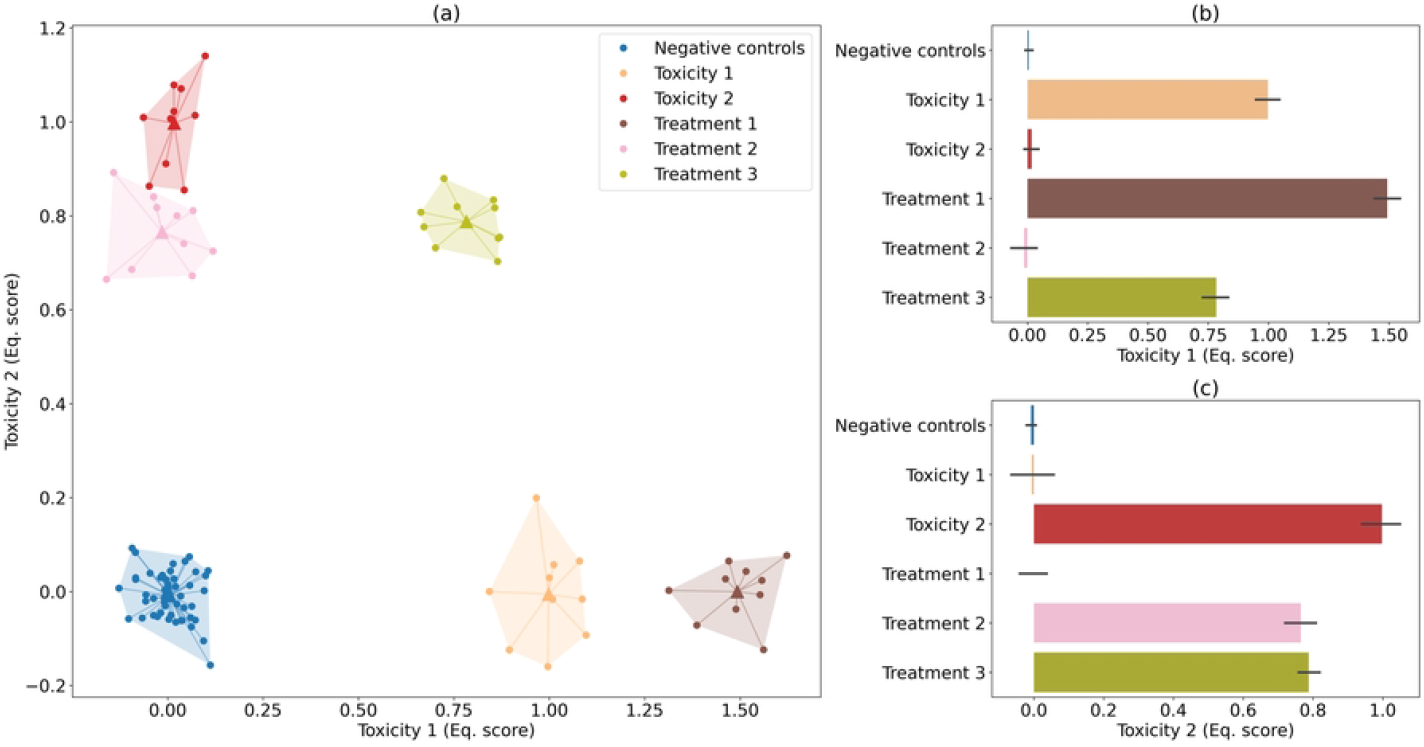
(a) A scatter plot showing how similar treatments are to each other in terms of the Eq. score of hypothetical Toxicity 1 and 2. On the x-axis, the Eq. scores from the model of Toxicity 1 are presented and vice versa for the y-axis and Toxicity 2. The observations of these groups are also present in the plot as orange and red clusters. (b) The Eq. scores of Toxicity 1 are presented as a bar plot. These are the same values used in (a) on the x-axis. (c) The Eq. scores of Toxicity 2 for different treatments.

By summarizing the feature vectors into Eq. scores and visualizing them in a scatter plot or bar plot, we can efficiently compare and interpret the toxic effects of different treatments. This approach offers a valuable tool for researchers in understanding the relationships between various treatments and their associated toxicities.

## Results

### Comparing Eq. scores

By fitting an PLS/OPLS model on a reference group of treatments and the negative control group and predicting the other compounds with it, we get Eq. scores. We can use the Eq. scores as axes in a scatter plot to see relations between treatments, such as in Fig. 4. In Fig. 4a we see distinct clustering of treatment groups. Noticeably, SB-203580 and SB-202190, which have the same target MAPK14, are close. Both have high Eq. scores of SB-202190 and low Eq. scores of purvalanol-a. We can observe the same with aminopurvalanol-a and purvalanol-a with the common target CDK2. These treatments instead have low Eq. scores of SB-202190 and high Eq. scores of purvalanol-a. This can also be visualized in bar plots as in (b) and (c) to see the comparison in one treatment direction.

**Fig 4.**
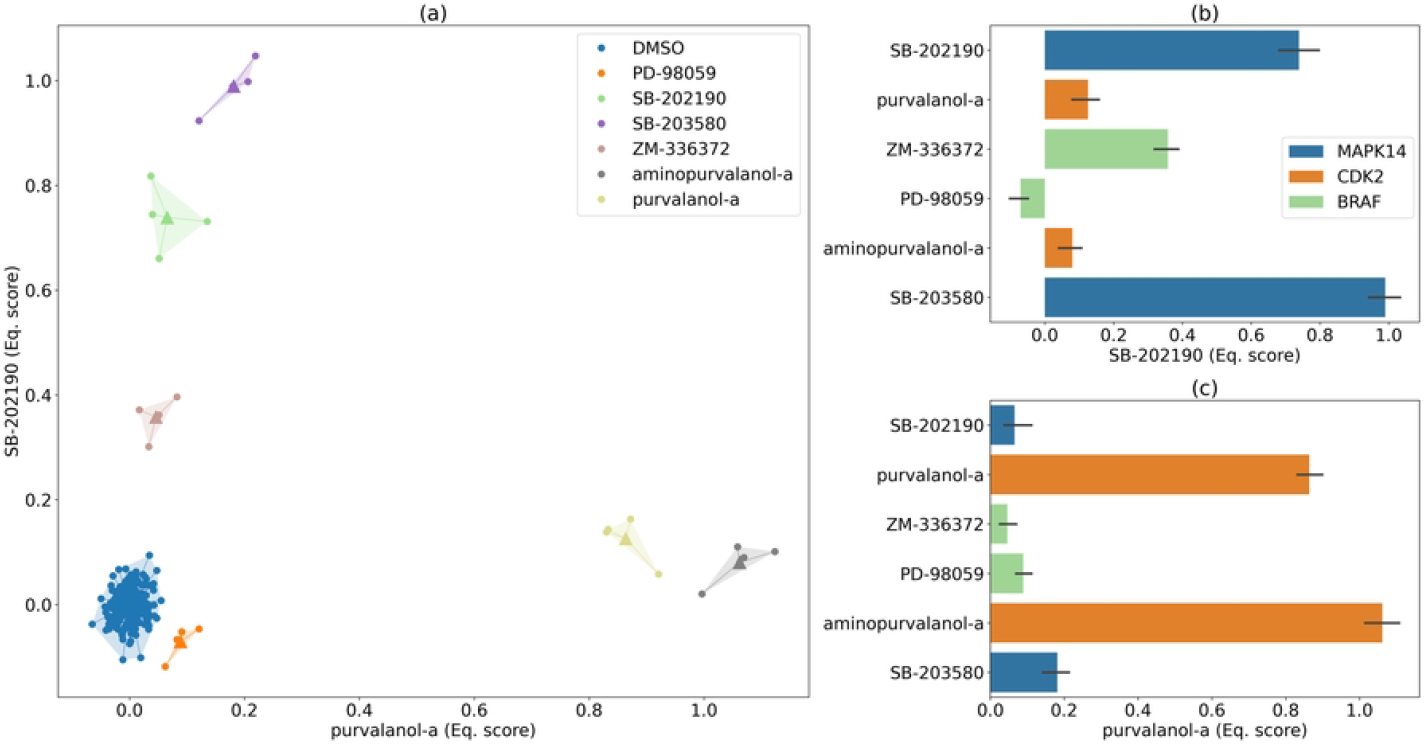
(a) A scatter plot showing how similar treatments are to each other in terms of the Eq. score of SB-202190 and purvalanol-a. (b) The Eq. scores of SB-202190 are presented as a bar plot. These are the same values used in (a) on the y-axis. (c) The Eq. scores of purvalanol-a for different compounds. As in (b) these values are also the same as in the x-axis (a).

Expanding further, we can incorporate the SSE of each model into the interpretation of the results. The SSE can also be used as a scaling factor to adjust and correct the predictions if they are not in a known feature space of the PLS model. Fig. 5 presents the results from models fitted on SB-202190, purvalanol-a and PD-98059 in bar plots. In Fig.5b, the Eq. scores from the SB-202190 model are displayed and we can see that SB-203580, which shares the same target, has the highest Eq. score. Furthermore, in Fig.5c it also has the lowest SSE, meaning that the prediction of the Eq. score is within the confidence space of the model. Based on these two scores, we calculate a weighted score in Fig.5a.

**Fig 5.**
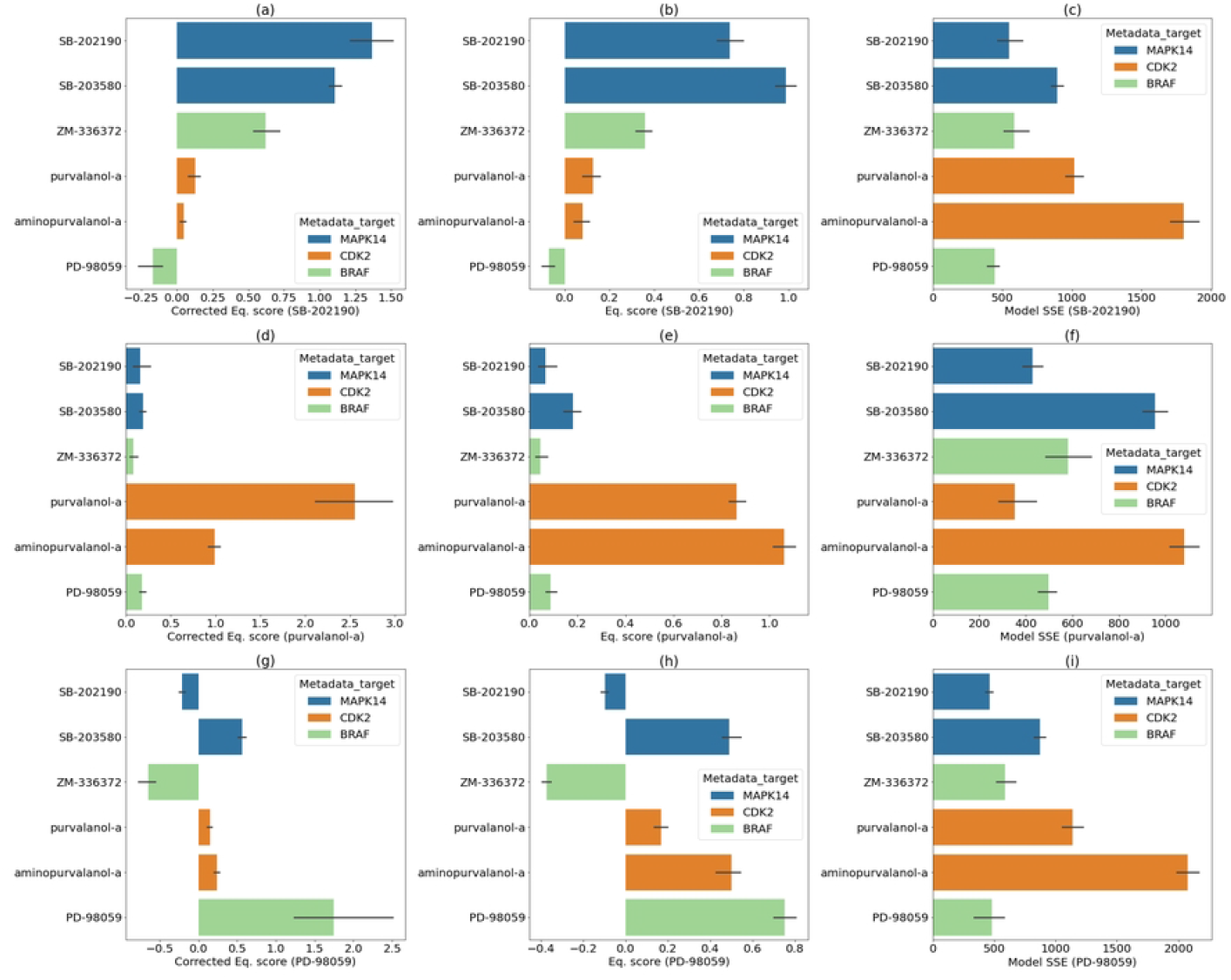
In the first row, (a), (b) and (c) the results from the model fitted on SB-203580 are presented. In the second row (d), (e), (f) the results from the purvalanol-a model and in the last row (g), (h), (i) are the results from the PD-98059. The bars correspond to treatments and are colored according to the treatment target where blue is MAPK14, orange is CDK2 and light green is BRAF.

Similar results can be observed in the second row for the purvalanol-a model’s prediction. However, since aminopurvalanol-a has a high SSE, the weighted score in Fig .5d is considerably lower than the Eq. score.

In the third row, we present the results from the PD-98059 model, and we observe a different trend. The treatment with the same target, ZM-336372, has a negative Eq. score. This indicates that the two treatments have an effect in the opposite direction with respect to the negative control group.

### PCA of Eq. scores

By combining the Eq. scores of the six compounds, we create a new feature space with the same number of features as compounds. By applying PCA to these new features we get principal components that capture the directions with the most systematic variation. A plot of the first two principal components is displayed in Fig. 6. The groups are distinctly separated. Notably, aminopurvalanol-a and purvalanol-a, which share the target CDK2, are both located in the same direction from the negative controls but with different magnitudes. PD-98059 and ZM-336372 with the common target BRAF, are located in different directions but close in T1. The opposite can be observed with SB-202190 and SB-203580 where the two are similar in T2 but not in T1. The same plot based on the original CellProfiler features can be found in the supplementary information.

**Fig 6.**
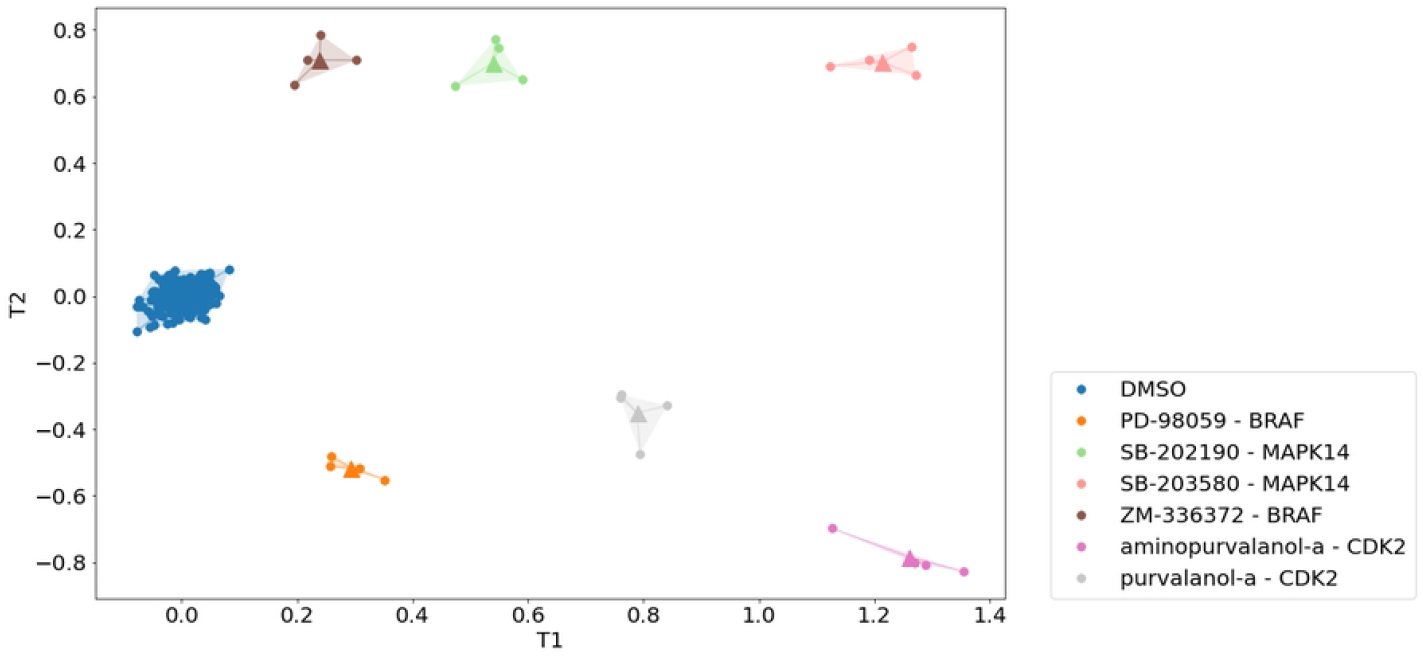
A PCA plot of Eq. scores from the compounds. T1 is the first principal component and T2 is the second principal component. There are four observations in each cluster and a center point marked with a triangle. DMSO is the negative control cluster located around the origo. In the legend, the name of the treatments and their corresponding targets are listed.

Fig. 6 summarizes the similarities of the compounds in a two-dimensional plane. It is important to note the difference between Fig. 6 and 5a. Fig. 5a uses the Eq. scores of two compounds as its axes while Fig. 6 uses the Eq. scores of all six compounds and projects into two dimensions.

### Benchmarking CellProfiler features vs Eq. Scores

So far, we have only looked at a portion of the full JUMP-CP1 dataset to comprehensibly illustrate the methods and its result. Now, we will apply it on a larger scale on all 303 compound treatments, cell lines and time points as in Fig. 2c. We use a benchmarking metric developed by Srinivas Niranj Chandrasekara at Broad Institute called the fraction of positive (fp) of the mean Average Precision (mAP) [14, 20]. This metric is presented in Fig. 7, for a comparison of replicability in terms of the fp between the predicted Eq. Scores, where the prediction is based on CellProfiler features, and the CellProfiler features themselves. Although the Eq. scores are based on the CellProfiler features, they perform consistently better than the CellProfiler features for the compounds due to the ability of PLS/OPLS models to capture and amplify structures in high-dimensional data.

**Fig 7.**
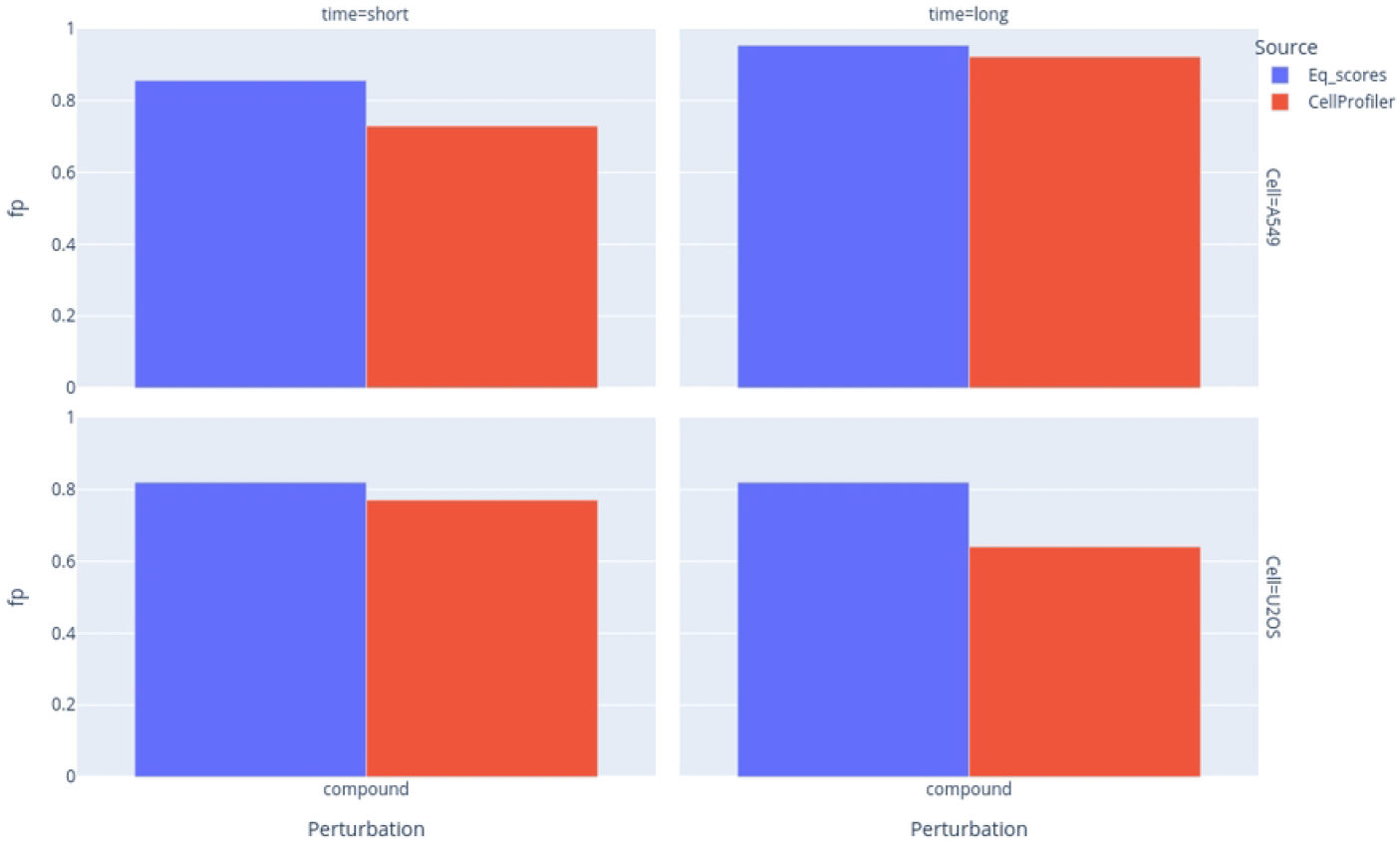
Comparison of the fraction of positive (fp) of the mean Average Precision (mAP) based on CellProfiler features (red) and the generated Eq. scores (blue) for compound treatments. The results are grouped according to cell type and at what timepoint the data was collected. Short represents the 48h mark and long the 96h mark.

In Fig. 8 we look at the mAP on which the fp in Fig. 7 are based, for the Eq. scores and CellProfiler features. Although the difference between the mAP of the Eq. scores and the CellProfiler features is not huge, the Eq. scores perform better across the board.

**Fig 8.**
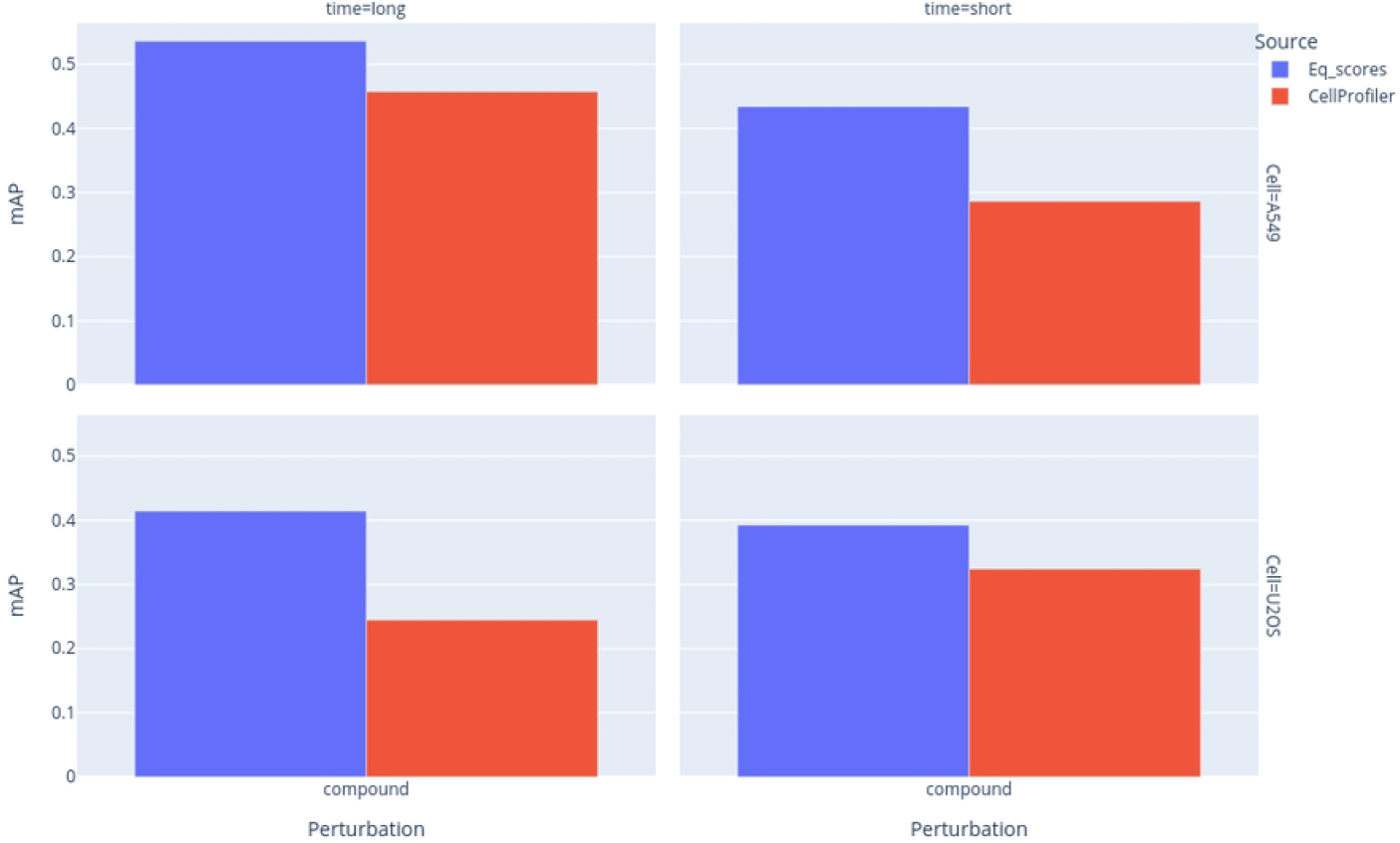
Comparison of the mean Average Precision based on CellProfiler features (red) and the Eq. scores (blue) for compound treatments. The results are grouped according to the cell type and at what time point the data was collected. Short represents the 48h mark whereas long represents the 96h mark.

## Discussion

Feature extraction from cell images and the post-processing of these are hot topics today, particularly within the biopharmaceutical industry. There are great tools available to extract these features but the tools to explore, interpret and discover similarities between different treatments, and especially between modalities, are falling behind. This is a challenging task since the number of features is generally high in comparison to the number of replicates. Luckily, this skewed relationship between observations and features has been the standard in chemometric problems for decades where the family of PLS methods has shown great success.

We have demonstrated that our proposed PLS/OPLS-based method can be useful for the downstream analysis of cellular morphological profile data. As shown with the compounds, we can compare treatments to one another by their Eq. score as well as using these scores as new features. The new features generated by the PLS/OPLS models represent the effect of each treatment in each treatment’s direction, defined by the PLS/OPLS models. These will summarize the hidden correlation of the most explanatory original features of each treatment.

The Eq. scores perform consistently better than the CellProfiler features in terms of the benchmarking metric defined by Niranj [20]. PLS/OPLS models focus on amplifying the features that are structurally important to distinguish a treatment from the negative controls. Thus, we should expect the Eq. scores to perform better than the original features as the models reconstruct the features resulting in a stronger, less noisy signal. Compounds typically have a clear effect in which the PLS/OPLS models can easily find the important variables and amplify the signal accordingly.

In drug screening and development, it is common that we know the gene we want to target to treat a disease but not which compound candidates target the correct genes. With this approach, we can more easily than before create an Eq. score of the gene or genes we want to target and then compare the equivalence of compounds and CRISPR-treated observations. Creating PLS/OPLS models based on the compound target and looking at the Eq. score of the CRISPR treatment will also give us more insight if the compound has other effects as well. This is a clear difference compared to correlation since correlations are symmetric and the PLS/OPLS models will be different.

We can also modify the method to suit different datasets. One example is where the concentrations of the compounds differ. In that case, the intensities set as the target for the PLS/OPLS models could be set as 1 and 2 for two different concentrations representing the different levels of effects. This, of course, is if the treatments are expected to behave in such a way.

We are looking forward to exploring the combination of chemometric approaches with the strengths of modern deep-learning approaches. We believe that these techniques can complement each other to increase our understanding of the complex data generated today and by doing so, decrease the cost and time to develop new, life-saving treatments.

## Acknowledgments

We would like to thank Pauala A. Marin Zapata at Bayer AG for her insights, curiosity and expertise in our discussions leading up to this work. The same goes for Srinivas Niranj Chandrasekaran, Anne Carpenter and Shantanu Singh at the Broad Institue for their support and never ending helpfulness.

